# Genome wide clustering on integrated chromatin states and Micro-C contacts reveals chromatin interaction signatures

**DOI:** 10.1101/2023.01.18.524458

**Authors:** Corinne E. Sexton, Mira V. Han

## Abstract

Chromatin states based on various histone modifications are a common annotation for genomes, and have been shown to correspond to regulatory functions such as enhancers and transcription start sites. With the advent of Hi-C and other chromatin conformation capture technologies, we now have the ability to analyze 3-dimensional physical interactions of chromatin regions, in addition to the 1-dimensional regulatory annotation, but methods to integrate this information are lacking. We present a framework for integrating the chromatin state of interacting regions into a numeric vector through the contact-weighted sum of chromatin states. Unsupervised clustering on integrated chromatin states and Micro-C contacts reveals common patterns of chromatin interaction which we call chromatin interaction signatures. Our results indicate that most chromatin interaction signatures are found in all four cell types investigated here. Between 12-40% of the regions change chromatin interaction signatures between the cell types despite maintaining chromatin state, hinting at the dynamic nature of chromatin conformation. Although regions with similar chromatin states are often in contact as expected, subcategories of enhancers and transcription start sites have distinct chromatin interaction signatures that are associated with gene expression. The integrated information on the chromatin states that the region is in contact with adds another layer of annotation beyond chromatin state or Hi-C contact separately. In summary, we present the first set of chromatin interaction signatures for the human genome that provides an integrated view of the complex dynamics of concurrent change occurring in chromatin state and in chromatin interaction.

## Introduction

Chromatin states are regulatory annotations defined by patterns of various histone modifications in genomic regions (Day et al. 2007). These patterns have elucidated regulatory roles of the noncoding genome (Hoffman et al. 2013). ChromHMM and Segway softwares use a Hidden Markov model and dynamic Bayesian network respectively to infer “hidden” chromatin states based on the combination of various epigenomic marks (Ernst and Kellis 2012; Chan et al. 2018; Ernst and Kellis 2017). This approach has been further extended to integrate additional molecular information in the model such as RNA-Seq, transcription factor binding and even 3D Hi-C data (Libbrecht et al. 2015; Wang et al. 2021; Liu et al. 2017; Mendez et al. 2020; Shokraneh et al. 2022). Each approach has in common the goal of annotating regions of the genome with interpretable clusters to be used in subsequent studies of gene regulation, genetic association, and others.

Of particular interest to this research is the integration of Hi-C data in the chromatin state inference framework. Hi-C sequencing captures the conformation of chromatin in the cell and this folding of DNA is an integral element of gene regulation. Though it is still of debate whether chromatin conformation is a cause or consequence of gene expression (Oudelaar and Higgs 2021), the canonical model assumes that distal enhancers bound by transcription factors require contact with promoters through DNA folding to influence transcription (Furlong and Levine 2018).

A challenge to integrating Hi-C in the previously stated chromatin state framework is the interacting nature of the data: a Hi-C contact involves two genomic regions, whereas chromatin mark ChIP-seq defines peaks in one region alone. To incorporate interaction into a feature space defined for each region, one has to summarize both the strength of the interaction that the region has with its multiple contacts, as well as the characteristics of each region in contact. A few softwares have attempted to integrate Hi-C in hidden state models in the following ways.

Segway-GBR (Libbrecht et al. 2015) and SPIN (Wang et al. 2021) both use integrative methods which encourage contacting regions to be clustered within the same state. This is based on the phenomena that large regions with similar chromatin marks are often in contact (Esposito et al. 2019; Hildebrand and Dekker 2020; Lieberman-Aiden et al. 2009). SPIN uses a Hidden Markov random field using a resolution of 25kb windows. Segway-GBR assigns a pairwise prior to significant Hi-C contacts at 10kb windows. Shokraneh et al (Shokraneh et al. 2022) employ graph embedding to learn structural vector features of 100kb resolution Hi-C data which were then passed on to an HMM. The resulting domains recapitulated known Hi-C sub-compartment categories (Rao et al. 2014).

Beyond these few examples, to our knowledge, no method exists to broadly investigate chromatin interactions and chromatin states across the entire genome. We propose a new method for integrating chromatin interaction data into a traditional framework for chromatin state annotation. Using this approach, we can apply genome-wide clustering to annotate patterns of chromatin interaction. By summarizing all contacts at each individual genomic region of interest, we capture a chromatin interaction signature (CIS) of both the summed strength of contacts as well as the chromatin states of those contacts. These CISs provide an additional layer of genome-wide annotation beyond contact or chromatin state alone. Applying this approach to Micro-C data (at 1kb window size) (Krietenstein et al. 2020; Hsieh et al. 2016) rather than lower resolution Hi-C, we were able to annotate chromatin interaction signatures for chromatin state segments that span smaller regions, such as enhancers or promoters, rather than looking at broad compartment activity.

## Results

### Genomic contacts are widespread across multiple unique chromatin states

Chromatin state as marked by distinct combinations of histone modification is directly related to the 3D folding of the genome (Boettiger et al. 2016). Active (A) and inactive (B) chromatin compartments characterized by Hi-C are enriched in corresponding active and inactive chromatin marks respectively (Rao et al. 2014). Interestingly, a recent paper also proposed an intermediate (I) state between A and B which is enriched for poised-promoter and polycomb-repressed chromatin states, marked mainly by presence of H3K27me3 marks (Vilarrasa-Blasi et al. 2021).

Though the interplay of several chromatin states within one compartment is well characterized, the impact of an individual genomic region in contact with several different chromatin states has not been studied extensively. To explore the global view of chromatin state interactions across different cell states, we focused on the four cell types (H1-hESC, HeLa, HFF, definitive endoderm) that have been recently assayed with Micro-C (Akgol Oksuz et al. 2021). Chromatin states used in this analysis correspond to the 18-state ROADMAP model inferred using ChromHMM (Kundaje et al. 2015) and were obtained for the four cell types from the EpiMap repository (Boix et al. 2021).

Important to note is that the distribution of chromatin states varies greatly between the four cell types (Figure 1). In particular, the endoderm cell type has many more quiescent characterized regions (those with low chromatin marks) compared to the other three cell types used here. Though this affects downstream analysis, we choose to include endoderm in our clustering because including a cell type with larger proportion of quiescent regions that have very low signals for all available histone marks shows that the method is robust to different distributions of marks.

**Figure 1.**
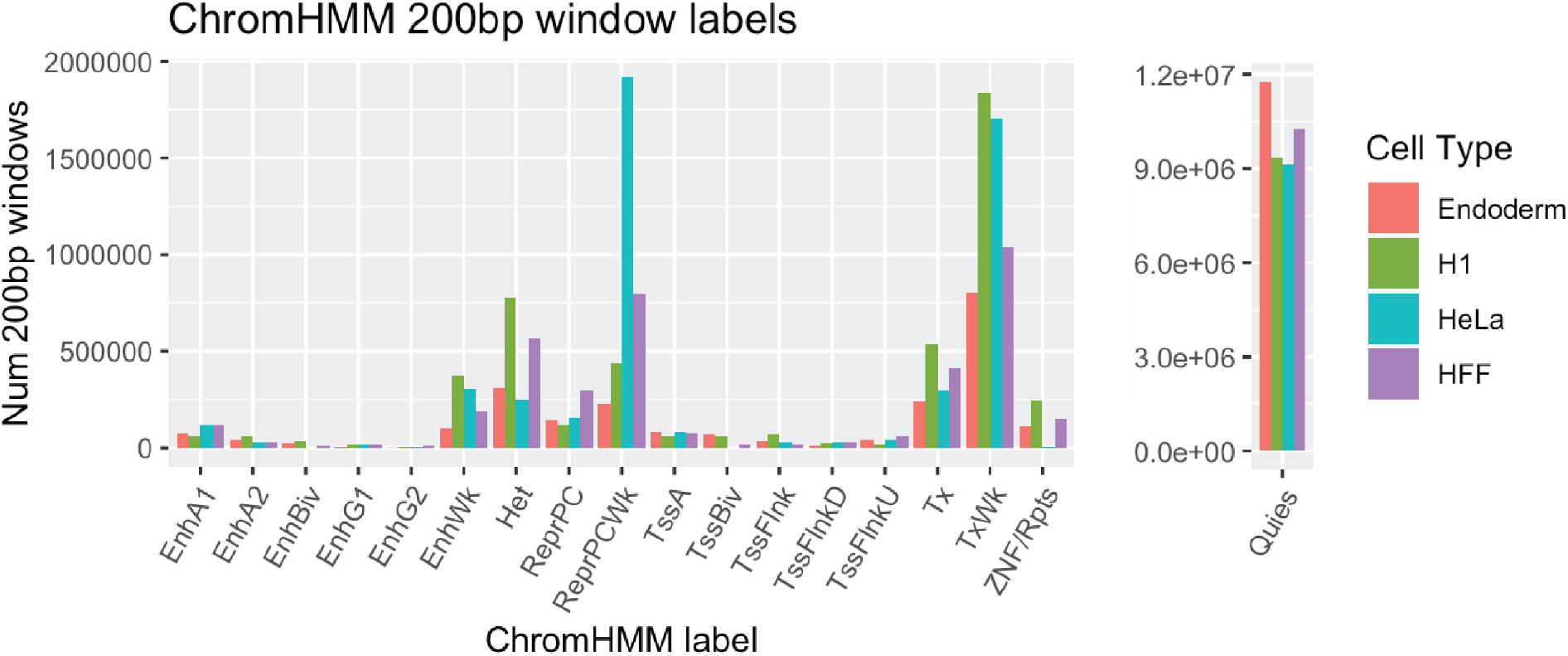
Chromatin state distribution between cell types. For endoderm, H1, HeLa, and HFF cells, annotated ChromHMM segments were divided into equidistant 200 bp windows to show the amount of ChromHMM state distribution between cell types.

To quantify the frequency of different chromatin state interactions for a single region, we counted the number of unique ChromHMM states that each genomic region is in contact with (Supplementary Figure 1). Here the region is defined by the segmentation of ChromHMM and thus can be of variable length. The median number of unique ChromHMM chromatin states in contact with any single region is 12 for endoderm and HFF cells, 11 for H1 and 13 for HeLa cells suggesting a high degree of connectivity between several different chromatin state types for any single region.

Importantly, this high degree of interaction is not due to the Hi-C window spanning multiple chromatin states, as one usually finds with window size of 5Kb or more. As we are examining micro-C interactions, we can observe the interactions between segments that are as small as 1 kilobase.

### Integrating chromatin interactions through the contact-weighted sum of chromatin states

Given the complex nature of interaction across different chromatin states, we need an approach that will summarize this high degree of interaction for any single region. We used a straightforward approach of summing across the different chromatin states, weighted by the contact intensity. For each region segmented with a chromHMM state annotation, all contacts 2Mb upstream and downstream were summarized by summing all KR-normalized Micro-C scores across corresponding chromatin states, as shown in Figure 2 (See Methods).

**Figure 2.**
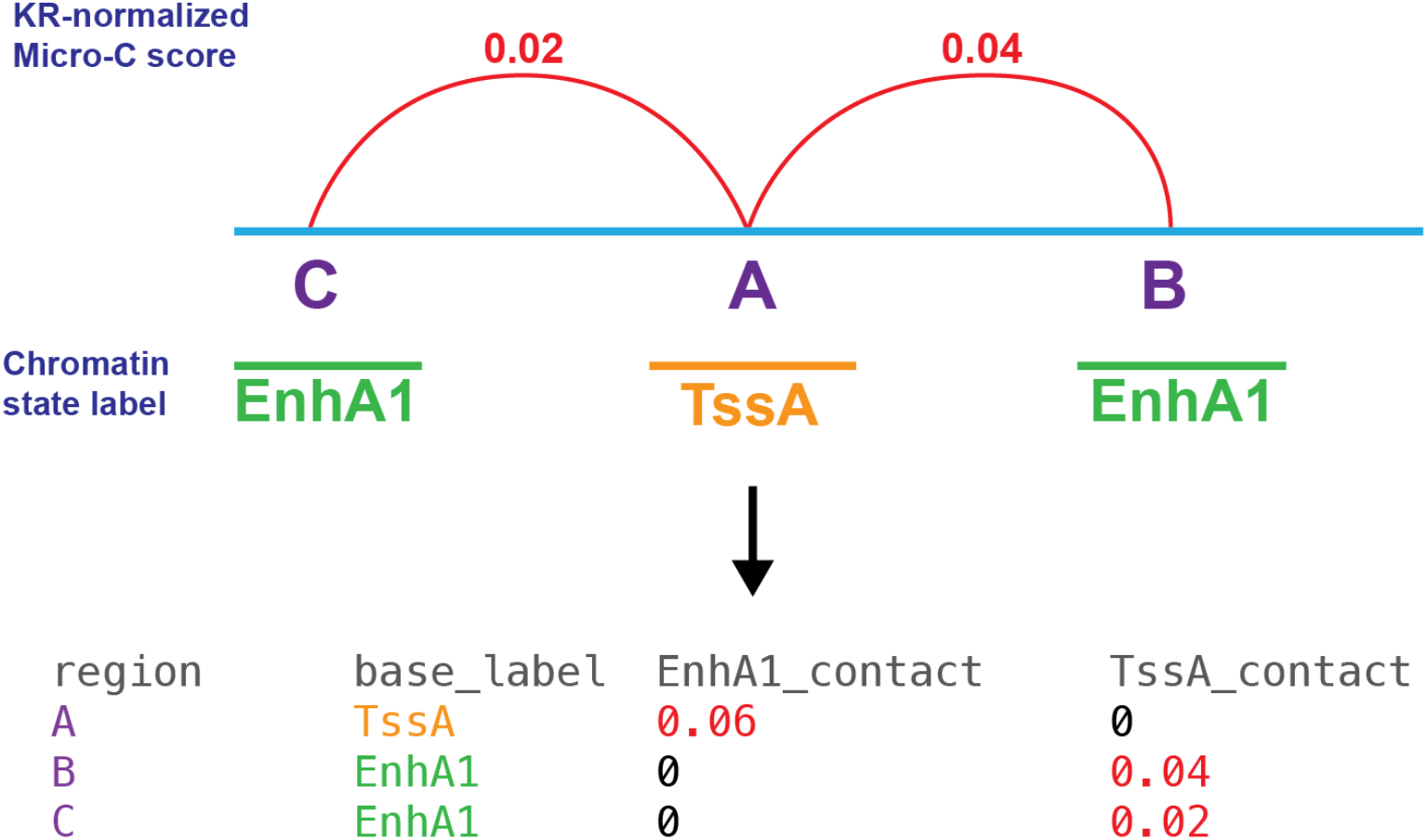
Creation of Sum of Chromatin state by Contact matrix. Region A is in contact with EnhA1 regions B and C and therefore the KR-normalized scores are summed for those regions and added to the matrix. Similarly, region B is in contact with region A and the KR-normalized score is included in the matrix for region B and likewise with region C.

Sum of Chromatin state by Contact (*SCC*) is defined as a *N* × *M* matrix, where *N* is the total number of segments annotated by ChromHMM and *M* is the number of possible chromatin states defined by ChromHMM. Each row of the matrix, *SCC_i_*, *i* = (1,…*N*), is defined for the focal segment *i*, which we call the base region. *SCC_i_* is a vector of length *M*, and is defined as the contact-weighted sum of all chromatin states that are interacting with segment *i*.

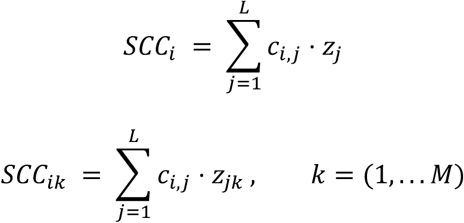

*z_j_* = (*z*_*j*1_,…*z_jM_*) is a vector that represents the chromatin state of the interacting segment *j* that is in contact with segment *i*. The vector consists of *M* binary indicator variables that are mutually exclusive and exhaustive (i.e., one and only one of the *z_jk_*’s is equal to 1, and the others are 0). This indicator vector is multiplied by the contact intensity *c_i,j_* between the chromatin segment *i*, and the interacting segment *j*. When the segment spans many Micro-C windows, the mean contact intensity is used. Then the contact weighted indicator vector is summed across all interacting segments *j* = (1,… *L*), within the ±2Mb window.

This approach assumes that each interaction with a chromatin state segment contributes additively and independently to the base region’s annotation. A similar additive approach was proposed for enhancer-target prediction in a model called Activity by Contact in Fulco et al. (Fulco et al. 2019). Our approach here is to extend the model to all chromatin states, rather than focusing on enhancers, as our goal is to identify broad patterns rather than specific enhancer-gene pairs.

As mentioned before, these regions were of variable length depending on the chromatin state segmentation. We chose to define base regions by chromatin state segmentation rather than a fixed window size because we are most interested in how annotated regions interact with other annotated regions. Having a fixed window size can be misleading in this regard, because due to our assumptions above, a large segment that is broken into multiple smaller windows would then be summed to result in a stronger value linear to the size of the segment, and we did not want that effect of segment size influencing our results. Also, further subdivision of chromatin states into smaller windows would not provide additional information to our clustering because the Micro-C resolution used was 1kb.

We plotted the resulting Sum of Chromatin state by Contact (SCC) matrix in Figure 3. We concatenated the matrices of all four cell types to detect patterns found across cell types. We excluded quiescent segments from the rows because patterns of contact with low marked regions are of little interest but being the most widespread segment in the genome (Figure 1), they occupy too many rows in the matrix if included. However, we included contacts to quiescent states in the columns to show the overall distribution of contact. A clear diagonal is evident which confirms the previously reported pattern that similar chromatin states interact with each other (Esposito et al. 2019; Hildebrand and Dekker 2020; Lieberman-Aiden et al. 2009). More interestingly, there are also off-diagonal hotspots suggesting that specific interactions between different chromatin states can happen frequently. Based on this observation, we employed unsupervised clustering to characterize the patterns of chromatin interactions for each region across the whole genome.

**Figure 3.**
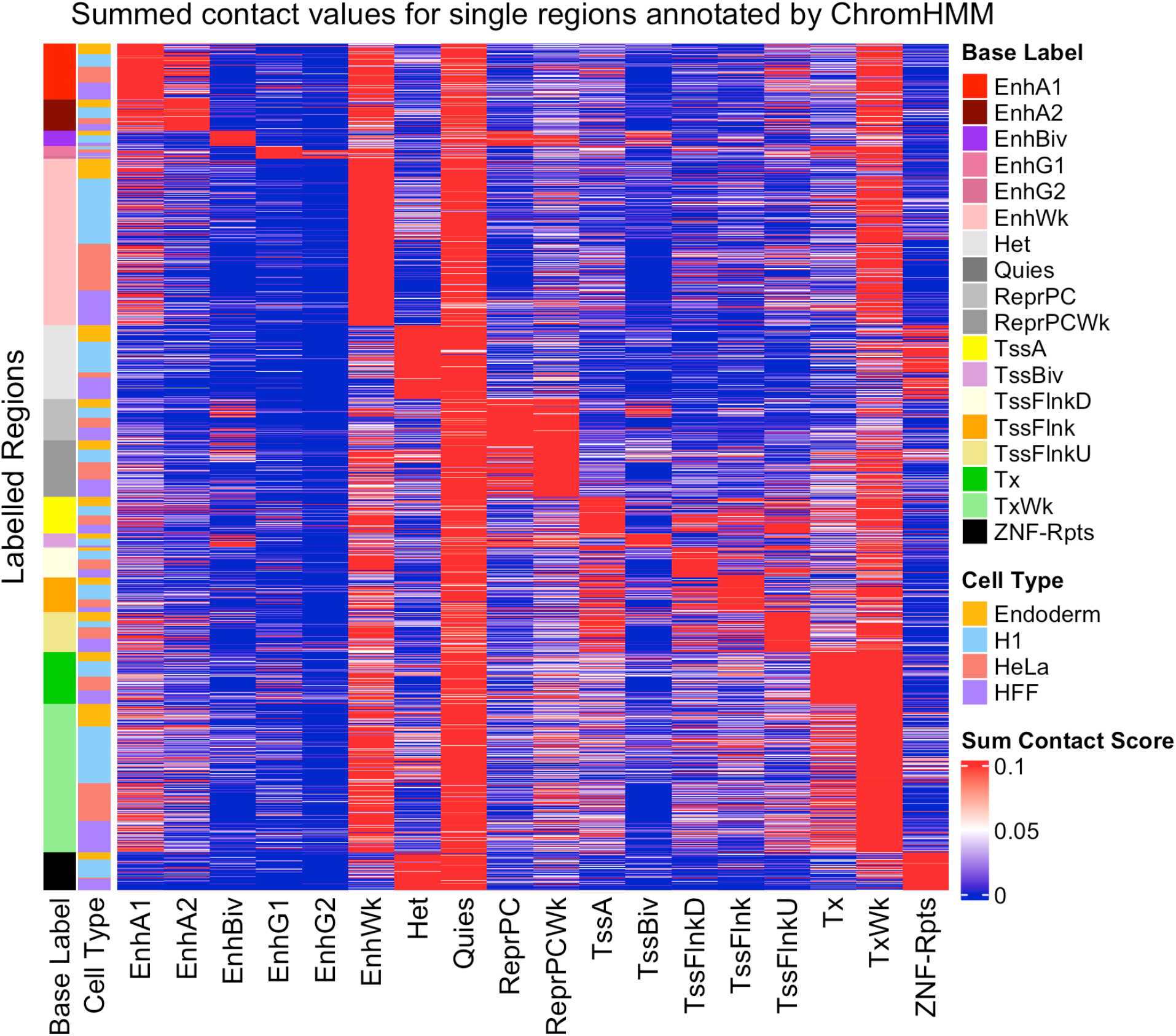
Sum of Chromatin state by Contact matrix. Each row is a ChromHMM annotated segment. Contact Score in each column represents the KR-normalized Micro-C contact scores summed across all the interacting segments annotated with the corresponding chromatin state. The heatmap is ordered by base chromatin state, then by cell type.

### Clustering on integrated chromatin states and Micro-C contacts reveals chromatin interaction signatures

The concatenated SCC matrix for all four cell types (Figure 3) was clustered using k-means and a k of 18 was chosen based on cluster interpretability and the “elbow” plot method (see Methods). While we included contact with quiescent regions in the columns of Figure 3, we excluded contacts with quiescent regions in our clustering analysis because they represent broad regions with low chromatin signals and can overwhelm the clustering. Results of k-means clustering (Figure 4) yielded several clusters with distinct chromatin interaction signatures (CIS). We coin this term to describe a distinct pattern of chromatin interactions for a single genomic region. After clustering, we named each CIS according to its pattern of contact. For example, regions in the cont_Tss_Enh cluster exhibit high contact with both transcription start sites (TSS) and enhancer regions. All 18 CIS names and their ratio of contact scores are shown in Figure 4.

**Figure 4.**
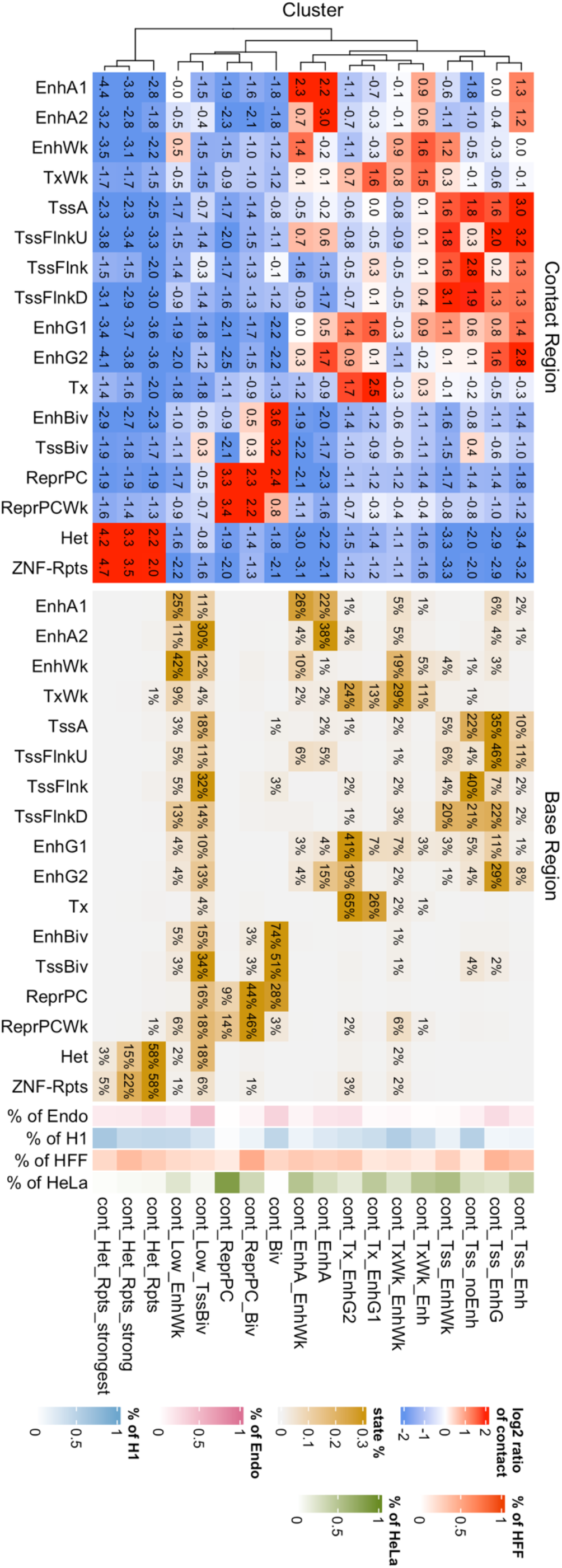
K-means clustering of the SCC matrix. Summed contact data for ChromHMM segmented regions (of variable length) for all four cell types were included in k-means clustering with a k= 18. The left panel shows the log2 ratio of mean summed contact scores for each segment divided by the mean summed contact scores across all clusters. In essence, this shows the enrichment of contacts to different chromatin states in each cluster. Included for annotation on the right is the percent of the base chromatin states found as well as the cell type amount present in each cluster.

The left panel of Figure 4 corresponds to the log2 ratio of the mean Micro-C score in contact with each chromatin state within a CIS cluster divided by the mean Micro-C scores in contact with each chromatin state across all CIS clusters. This shows the enrichment of each contact type for all 18 CIS clusters. The right panel is for annotation purposes only and was not included in clustering. The percentages correspond to the percent of the chromatin state of the base regions which belong to each CIS cluster, with each column summing to 100%. Lastly, the rightmost cell type annotation shows what percent of the cluster is made of regions from each cell type. For example, the cont_ReprPC cluster is mostly made of HeLa cell type regions.

Several clear patterns emerge. The first notable pattern is that across all regions with various base ChromHMM states, a subset of the regions are grouped into the Low contact clusters (cont_Low_TssBiv and cont_Low_EnhWk), while the rest are distributed across various clusters of high contact, showing clear difference in the contact intensity even among regions with the same chromatin state annotation.

The second pattern is that the enriched interaction between the same chromatin states that showed up as the high scores along the diagonal in the heatmap (Figure 3), are also present as enriched concentrations in Figure 4. For example, a distinct signature of contact with bivalent states for bivalent enhancers and bivalent TSS, and enriched contact with repressed polycomb states for repressed polycomb regions are clearly notable.

The third pattern we see is that regions with the same base chromatin annotations are now subdivided into different clusters based on the chromatin states they are interacting with. For example, the cont_Tss_Enh cluster is contrasted with the cont_Tss_noEnh cluster, the latter of which is depleted of enhancer contact, but is highly enriched for TSS contact. If one focuses on the base region TssA column (Active TSS), we can see that 35% of TssA regions cluster into cont_Tss_EnhG, but a sizable minority also group into cont_Tss_noEnh or cont_Tss_Enh clusters. Likewise for EnhG2, 29% of these regions cluster into cont_Tss_EnhG, but less often it also groups into cont_Tss_Enh cluster. This perhaps shows a subdivision of enhancers with some elements preferring singular contact with TSS and others interacting with enhancers as well.

### Each CIS is present in all cell types, but individual regions often change CIS between cell types

To understand how the genome-wide chromatin interaction changes across cell types, we compared the CIS clusters across cell types. Four cell types were included in this analysis: H1-hESC, HeLa, HFF, and definitive endoderm. These cell types were chosen based on the availability of human Micro-C data (Akgol Oksuz et al. 2021). Each of the cell types have regions with membership in all 18 CISs, hinting at the common pattern of chromatin interaction even among diverse cell types. This is similar to how A and B chromatin conformation compartments are observed across cell types.

A notable exception is the cont_Biv cluster which has much less HeLa membership than the other cell types (Figure 4). This is most likely due to the lack of TssBiv and EnhBiv chromatin state regions in HeLa cells (Figure 1). Similarly, HeLa cells also have a lower proportion of Het regions and therefore have less membership in the 3 cont_Het_Rpts clusters.

Though the same CISs are consistently found in all four cell types, individual regions often change CIS between cell types (Figure 5). Of particular interest are regions where chromatin state stays constant, but CIS is different between cell types. Depending on cell type comparison, between 12%-40% of 200bp base regions cluster into a different CIS despite retaining the same chromatin state between cell types (Figure S3). This suggests that contact itself may indeed provide another layer of annotation beyond chromatin state. We chose to use a fixed window size of 200bp in this visualization, because it is the smallest unit of the chromatin state segmentation observed in the data, and shows more accurately the distribution of chromatin state and CIS changes between the four cell types which each have varying window sizes due to the nature of cell type specific chromatin state segmentation.

**Figure 5.**
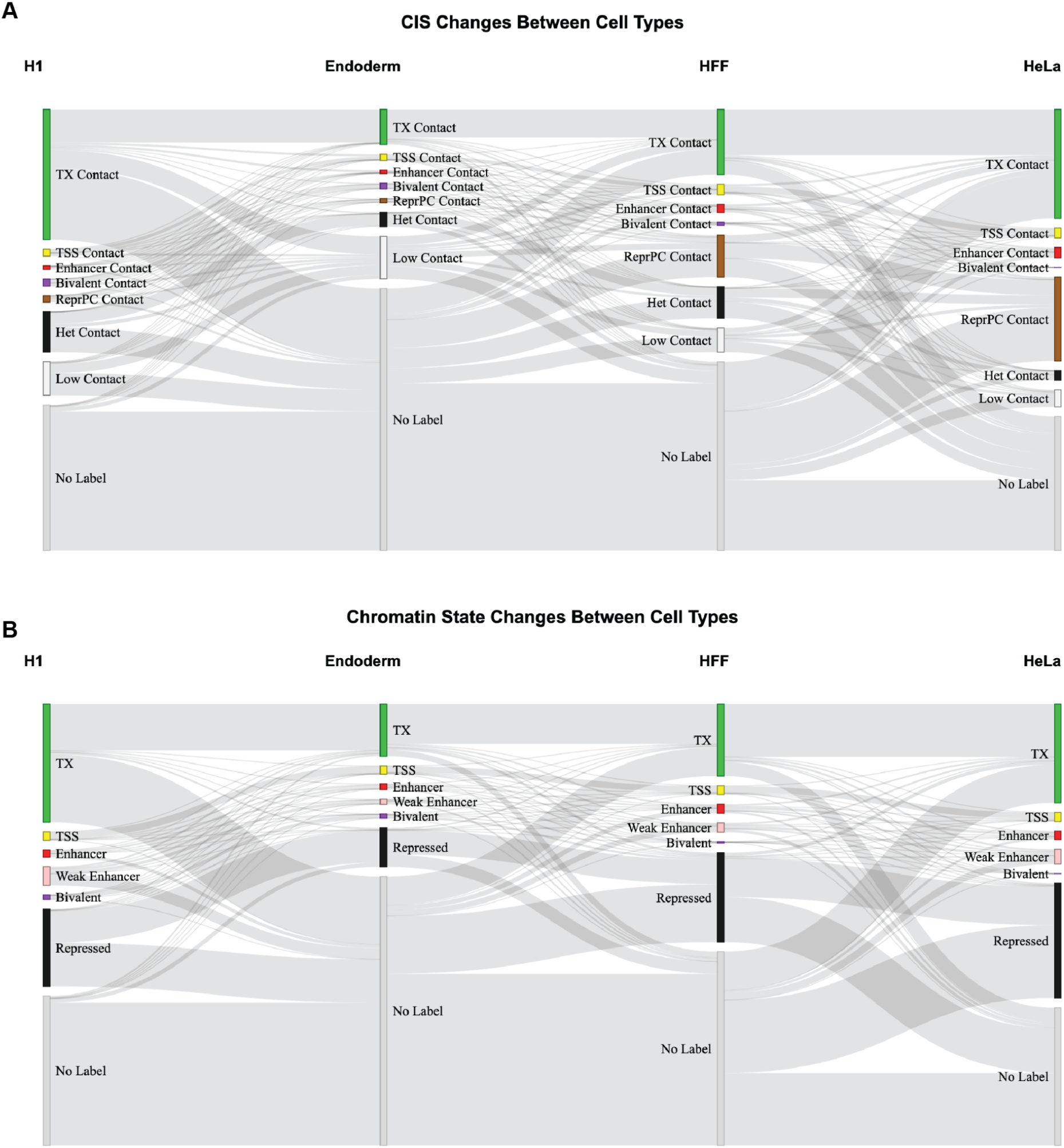
CIS and Chromatin State Changes Between Cell Types. Sankey diagrams show the change of CIS (A) and chromatin state (B) for 200bp regions between H1, endoderm, HFF, and HeLa cell types.

### Changes in CIS in the transcription start sites are associated with change in gene expression

To assess functional significance of CIS, we investigated CIS clusters at the transcription start site (TSS) of upregulated and downregulated genes (Figure 6). We also looked at all TSS regions in general (Figure S3).

**Figure 6.**
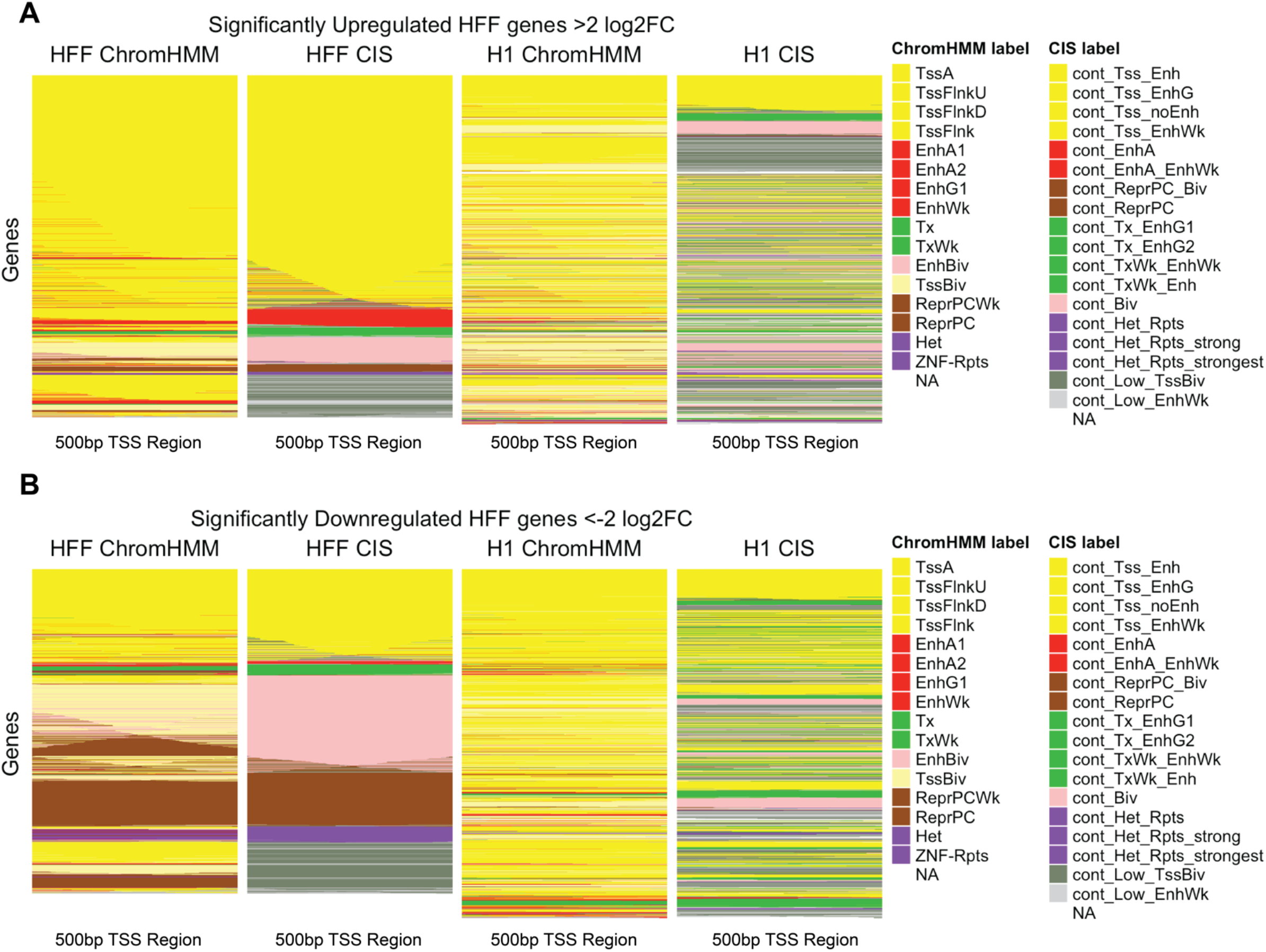
CIS and chromatin states for differentially expressed genes. Genes with adjusted p-value less than 1e-10 and with an HFF vs H1 log2 fold change >2 for upregulated genes (A) and <-2 for downregulated genes (B) were selected between H1 and HFF cell types. CIS and chromatin states are shown for each gene’s 500bp TSS region. The genes (heatmap rows) are ordered by proportion of clusters in the order

We looked at genes with greater than 2 log2-fold gene expression change in HFF vs H1 cell types (Figure 6). In general, TSS chromatin states are TssA, TssFlnkU, TssFlnkD, or TssFlnk for upregulated genes and TssBiv and ReprPC for downregulated genes as expected. However CIS are more variable and therefore provide additional information beyond chromatin marks alone. The TSS of genes upregulated in HFF (Figure 6A) show increased proportions of cont_TSS, cont_EnhA, cont_Tx and the TSS of genes downregulated in HFF (Figure 6B) show increased proportions of cont_Biv, cont_Het, cont_ReprPC and cont_Low. A similar pattern can be seen in H1, when the regions are ordered by H1 CIS as well, with specific CIS being more pronounced (Figure S4). To understand the information gain provided by CIS, we quantified the normalized mutual information (NMI) between the chromatin state and CIS change in the TSS and the gene expression change, comparing HFF and H1. The NMI between gene expression (up or downregulation) and chromatin state change was 0.19, and the NMI between gene expression and CIS change was 0.15. While CIS shows less amount of information dependency with gene expression, it is notable that without knowledge of the chromatin state of the TSS itself, one can gain about three quarters of information by observing the chromatin states of interacting partners, as by observing the chromatin state of the TSS directly. When we concatenate the CIS and ChromHMM states, the NMI is 0.21, showing we gain information by observing CIS together with chromatin state, compared to just observing the chromatin alone.

We looked at the well-characterized H1 pluripotency gene *NANOG* (Boyer et al. 2005), to understand the utility of CIS clustering compared to chromatin state alone. We show a detailed example of changes between H1 and endoderm cells for the *NANOG* TSS region (Figure 7).

**Figure 7.**
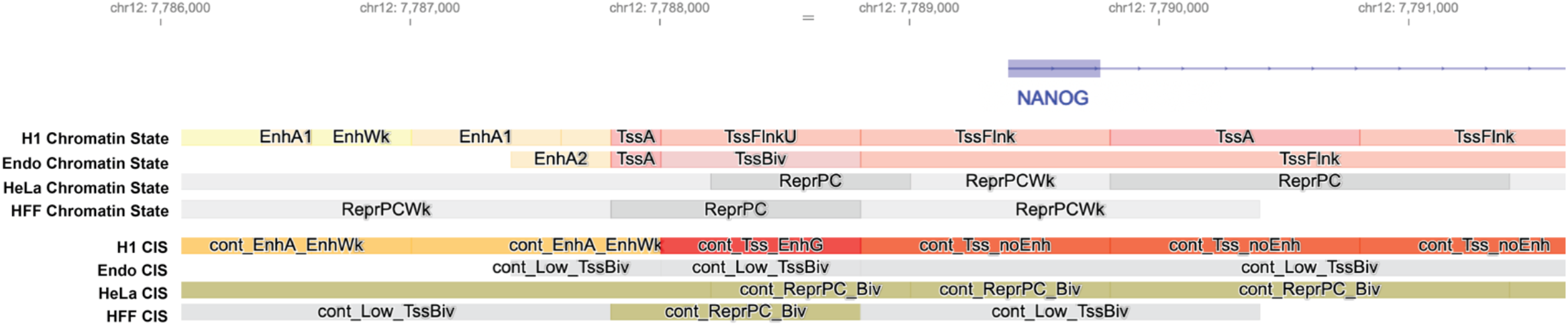
Chromatin state and CIS annotations at the *NANOG* TSS. Though the endoderm and H1 chromatin state annotations are largely similar, the CIS shows extensive differences. This corresponds with decreasing NANOG expression as H1 develops into endoderm cells.

The chromatin state is largely identical in both endoderm and H1 cells, except for a change from TSSFlnkU to TSSBiv for a small region near the promoter, consistent with the pattern observed for TSS regions in Figure 6. However, the CIS shows more extensive change between the two cell types, with endoderm cells clustered as Low contact and H1 cells clustered as Enhancer contacting. In this case, the CIS adds extra information which correlates with the increased *NANOG* expression found in H1 cells.

Understanding the reasons behind the CIS change are crucial as clustering is performed on a summed contact matrix. What is the main driver for the difference between endoderm and H1 CIS? Because of the way that we define SCC as a summation vector for each region (Figure 2), 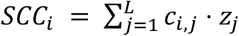, there are only two ways that can result in change in this vector. Either the strengths of the contacts can change (*c_i,j_*), or the chromatin states of the interacting regions can change (*z_j_*). We decided to look at the TSS of *NANOG* in detail, to understand the components driving the change in CIS. The details of the chromatin interactions and the chromatin state of the interacting regions are shown for the *NANOG* TSS in Figure 8.

**Figure 8.**
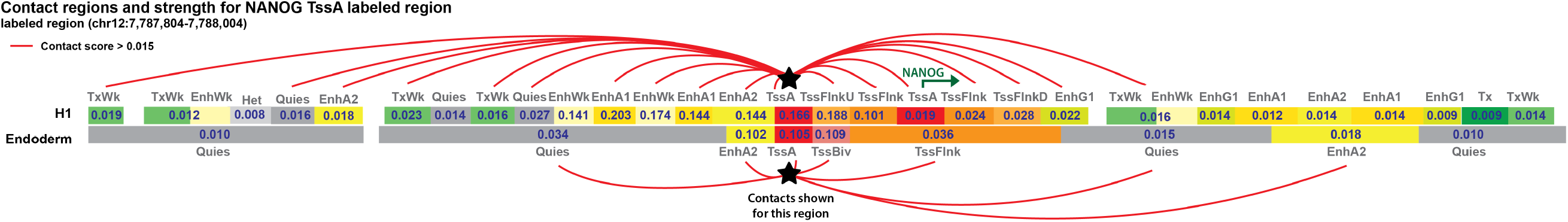
Bars are representative of genomics regions in contact with *NANOG*. They are in genomic order, but are not to scale. Regions are labeled by chromatin state and weighted Micro-C strength. Red connections signify Micro-C contact with a KR-normalized score above 0.015.

In this example, we observe a dynamic change in both contact strength and chromatin state for most regions in contact with the TSS. This is to be expected as chromatin interactions are known to occur between similar regions and therefore a change in histone marks can affect contact strength. There are however a few regions in this example which maintain chromatin state such as the TssFlnk regions found around the *NANOG* TSS in both cell types and the EnhA2 region downstream of the *NANOG* TSS. This example shows how our method is able to integrate the complex dynamics of concurrent change in chromatin state and in chromatin contact and summarize it down to lower interpretable dimensions. The CIS annotations that represent the combination of chromatin state and Micro-C data make it easier to identify regions of interest that are undergoing change in both chromatin interaction and in the chromatin state of its interacting regions. This is a unique strength of our approach, since such identification is not possible based on chromatin state or Micro-C data alone.

### Active enhancers and super enhancers are enriched in CIS clusters

To determine if unsupervised clusters are enriched in cell-type specific regulatory elements, we conducted enrichment analyses looking at relevant enhancer and promoter regions in each cell type. For each cell type, we conducted chi-squared tests within each cluster for super enhancer annotations (Jiang et al. 2019) and FANTOM5 CAGE-defined active enhancers (Andersson et al. 2014) as well as TSS regions for highly expressed genes and lowly expressed genes (see Methods).

As shown in Figure 9, several clusters show significant and strong enrichment across cell types for unique annotations. Of particular interest are the cont_Tss_Enh and cont_Tss_EnhG clusters which are enriched in active enhancers in both H1 and HeLa cells. These clusters are marked by strong association with TSS and other enhancer regions.

**Figure 9.**
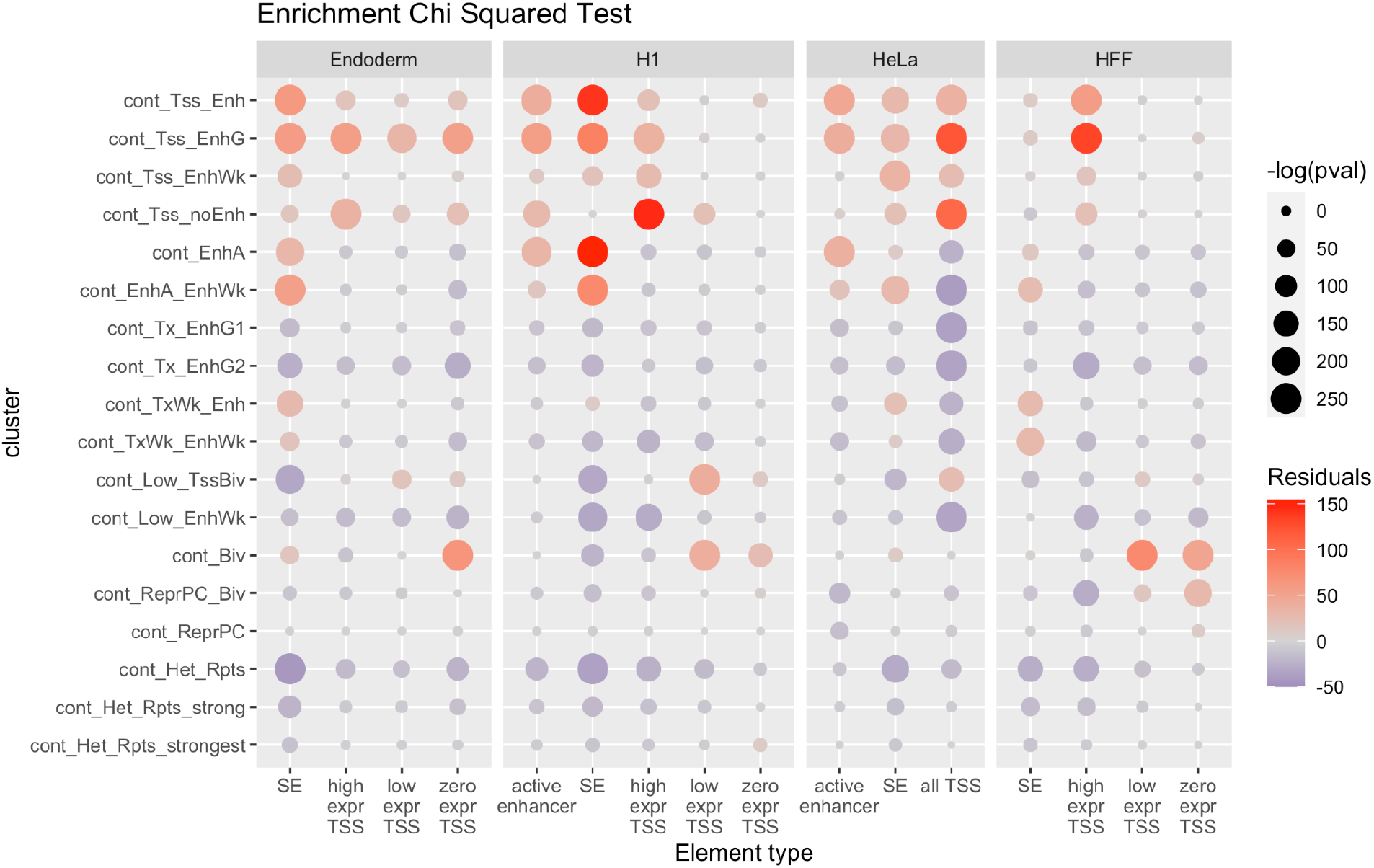
Enrichment results for CIS of all four cell types. Based on data availability, we performed chi-squared tests to test enrichment for super enhancers, high, low, and zero gene expression TSSs and active enhancers in each CIS cluster.

Regions clustered into cont_Tss_noEnh are enriched in the TSS of highly expressed genes in each cell type and less enriched for active enhancer regions. Cluster cont_Tss_noEnh represent regions that are mainly in contact with TSS associated chromatin states but not in contact with enhancer associated chromatin states.

Another interesting pattern is super enhancer enrichment in enhancer-contacting clusters across all cell types. Clusters cont_EnhA and cont_EnhA_EnhWk are associated with strong enhancer contact and weak TSS contact which is to be expected as super enhancers are generally defined as clusters of active enhancers (Hnisz et al. 2013).

A last observation is the enrichment of cont_Biv regions in lowly and zero expressed genes. Bivalent marked regions are purported to be “poised” regions of the genome marked simultaneously by both active and inactive chromatin marks (Bernstein et al. 2006). Therefore, the enrichment of low expressing TSS regions in the cont_Biv cluster could support the switching on/off of genes.

## Discussion

Methods for integrating 3D Hi-C data and traditional 1D data such as ChIP-Seq are a crucial step towards understanding the shifting dynamics of cell regulation. We show that clustering on an integrated matrix of Micro-C scores and chromatin states adds an additional layer of annotation to the genome.

Though chromatin states are known to interact with similar chromatin states in the genome (Esposito et al. 2019; Hildebrand and Dekker 2020; Lieberman-Aiden et al. 2009), we show that there are regions in contact with different chromatin states. These associations present an opportunity for pattern detection which can further aid with interpretation.

As shown in our enrichment analysis, cont_EnhA and cont_EnhA_EnhWk clusters are enriched for super enhancers, cont_Tss_Enh and cont_Tss_EnhG are enriched for active enhancers. Therefore, by clustering across the entire genome, we can effectively narrow the list of candidate regions with regulatory potential based on both contact and chromatin state evidence.

Most importantly, we present a framework for integrating chromatin conformation data into a more traditional vector format across the genome. This creates possibilities for future machine learning and clustering approaches. Additionally, this summative approach could be extended to include ATAC-seq, transcription factor binding, or DNA methylation data to further enhance regulatory predictions.

One caveat to our approach is the loss of information on specific pairwise interactions as a result of summing across all interactions with the same chromatin states. Future work to remedy this could include clustering based on pairwise relationships rather than on a summarized vector of contacts for one each base region. An alternative approach would be to utilize the graph structure of the contact data in order to preserve the individual contacts, and use graph embedding techniques to transform relational data into a vector form (Karbalayghareh et al. 2022). This would allow one to learn on genome segments (nodes) as well as pairwise contacts (edges), and has been shown to be a powerful approach.

In conclusion, we show a simple and straight-forward methodical approach to integrate contact and chromatin mark data across the genome, allowing researchers to distill complex chromatin interaction information into a vector, and then to an interpretable annotation, by further clustering on the vector. Using this method, we present a set of chromatin interaction signatures (CIS) that are frequently and repeatedly found across the genome and cell types. We also show that these chromatin interaction signatures allow a genome-wide view of chromatin interaction, and provide more information about gene regulation than either chromatin state or Hi-C contacts alone.

## Methods

### Data Sources

Chromatin states generated by ChromHMM were obtained from EpiMap for HFF, endoderm, H1-hESC, and HeLa cells for hg38 (Boix et al. 2021). The 18-state Roadmap model was used for these annotations, generated from H3K27ac, H3K4me1, H3K4me3, H3K36me3, H3K9me3, and H3K27me3 chromatin marks (Kundaje et al. 2015).

Micro-C mcool files were downloaded from the 4dnucleome.org and were originally from Oksuz et al (Akgol Oksuz et al. 2021). The Formaldehyde+DSG protocol was used for each of these Micro-C datasets. The HeLa cell data used was non synchronized as opposed to the cell cycle synchronized HeLa Micro-C data available.

A common issue with integrating chromatin mark data and Hi-C is the issue of window size. In general Hi-C data is interpreted in greater than 5kb windows whereas ChIP-Seq peaks are much smaller. For example, the smallest ChromHMM segmented regions are 200bp. To remedy this, we take advantage of available Micro-C data. Micro-C data is a derivative of Hi-C which uses a micrococcal nuclease (MNase) to create much smaller windows sizes (Hsieh et al. 2015, 2016). These high resolution fragments accurately capture fine scale interactions rather than the broader scale of traditional Hi-C enabling us to use 1kb windows for contact data.

### Creation of Data Matrix

Through a process we named Sum of Chromatin state by Contact (SCC), we create a matrix of Micro-C scores summed by chromatin state. The mathematical definition of the SCC matrix has been described in the results, here we describe the algorithm. Using Cooler (Abdennur and Mirny 2020), all Micro-C normalized weighted contacts were pulled 2Mb upstream and downstream for each non-overlapping 1kb window which contained an annotated chromatin state (omitting quiescent regions for feasibility purposes). Next, for each base region with a chromatin state, all regions in contact with the base were assigned a chromatin state based on results from bedtools intersect between ChromHMM segments and Micro-C bedpe files. In 1kb windows where multiple chromatin states are present, each is included as a separate entry. If a ChromHMM segment overlapped multiple windows, the mean Micro-C contact score across the windows was used.

Finally, the scores are summed for each unique chromatin state, resulting in a matrix where each chromatin state annotated base region (row) has 17 scores (columns), one for each chromatin state except for quiescent regions. Quiescent regions were excluded for this analysis because they signify regions with low chromatin marks. For clarity, the score is the summed amount of KR normalized contact the base region has with a particular chromatin state annotated region. All code for this process can be found at https://github.com/HanLabUNLV/SCC.

### K-means Clustering

After creating the matrix, *k*-means clustering was performed. To assess an appropriate *k* value of clusters, we employ the elbow plot method (Marutho et al. 2018). In brief, the sum of squared error for each cluster is calculated for several *k* values and the “elbow” point of the resulting plot is used to determine an appropriate *k* value (Supplemental Figure 2). We chose *k* = 18 based on the elbow method and to maximize understandability.

### Differential Expression and CIS Association

Bulk RNA-Seq count data from Chu et al (Chu et al. 2016) (GSE75748) was analyzed with DESeq2 to obtain differentially expressed genes between HFF and H1 cells. Up regulation was determined by >2 log2 fold change and down regulation was determined by <-2 log2 fold change and significance was detected at adjusted p-value < 1e-10. TSS sites were obtained from Fulco et al (Fulco et al. 2019) which narrowed the scope by selecting the single TSS for each gene with the largest number of coding isoforms. Bedtools intersect was then used to overlap those TSS regions with CIS and chromatin state annotations.

Specifically, every CIS-defined region for each cell type was intersected with the TSS regions. Then the different cell types were merged based on the genes they overlapped. These regions were then used for analysis of CIS changes (Figure 5). The 18-CIS model was collapsed to 7 labels for ease of interpretation. The membership of the 7 labels is as follows:

TX Contact: cont_Tx_EnhG1, cont_Tx_EnhG2, cont_TxWk_Enh, cont_TxWk_EnhWk
TSS Contact: cont_Tss_Enh, cont_Tss_EnhG, cont_Tss_noEnh, cont_Tss_EnhWk
Enhancer Contact: cont_EnhA, cont_EnhA_EnhWk
Bivalent Contact: cont_Biv
ReprPC Contact: cont_ReprPC_Biv, cont_ReprPC
Het Contact: cont_Het_Rpts, cont_Het_Rpts_strong, cont_Het_Rpts_strongest
Low Contact: cont_Low_TssBiv, cont_Low_EnhWk

#### Mutual Information Analysis

To quantify mutual information between differential gene expression (up or down regulation) and differential CIS/chromatin states, we concatenated the states for the HFF and H1 cell types. We used the aricode package (Chiquet et al.) to calculate the normalized mutual information using the NMI function for three different variables. i) NMI between chromatin states and gene expression ii) NMI between CIS and gene expression, and iii) NMI between the concatenated vector of chromatin state and CIS and gene expression.

### Enrichment Analyses

#### Data sources

To determine functional significance of the clusters, we calculated the enrichment statistics of enhancer and promoter elements for each cluster. FANTOM5 active CAGE-defined enhancers were obtained for each of the four cell types (HFF, HeLa, H1-hESC, and Definitive endoderm) (Andersson et al. 2014). Super enhancer annotations were taken from SEdb (Jiang et al. 2019). All TSS for the hg38 genome were obtained from refTSS (Abugessaisa et al. 2019) and high vs low expression genes were determined by taking the top 25% and bottom 25% of normalized gene expression counts for the following datasets: H1-GSE102311, Endoderm (Blake et al. 2018)-Additional File 1:Table S8C, HFF-GSM2448852.

#### Chi-Squared tests

To determine significance of enrichment, chi-squared tests were performed for each cluster within each cell type for each of the five sub groupings: active enhancers, super enhancers, high expressed gene TSS, low expressed gene TSS, and zero expressed gene TSS. Resulting p-values were Bonferroni corrected.

## Data Access

All code for the SCC method can be found at https://github.com/HanLabUNLV/SCC.

## Acknowledgements

This work was supported by the National Science Foundation under Grant No. 1750532 and No. 1946082.

## Supplementary Figures

**Figure S1.**
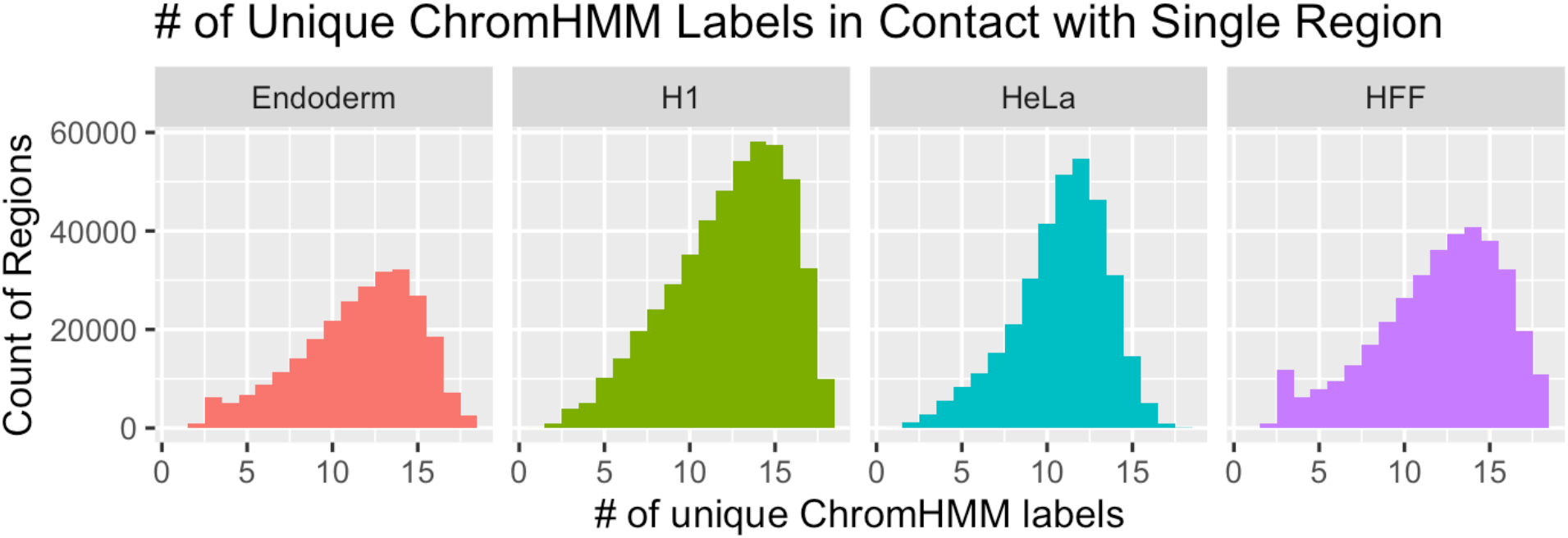
Number of unique ChromHMM states in contact with a single region. We counted the number of unique ChromHMM states that each genomic region is in contact with, with a non-zero micro-C score, for each cell type.

**Figure S2.**
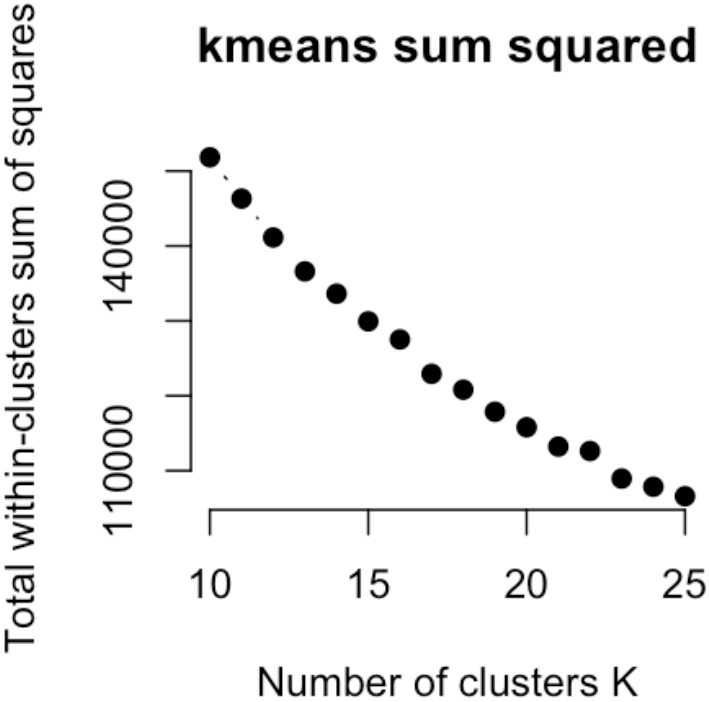
Elbow plot of sum squares calculations for k-means. To determine the number of clusters *k* to be used in our analysis, we employed the elbow plot method and looked for the most interpretable clusters (k = 18)

**Figure S3.**
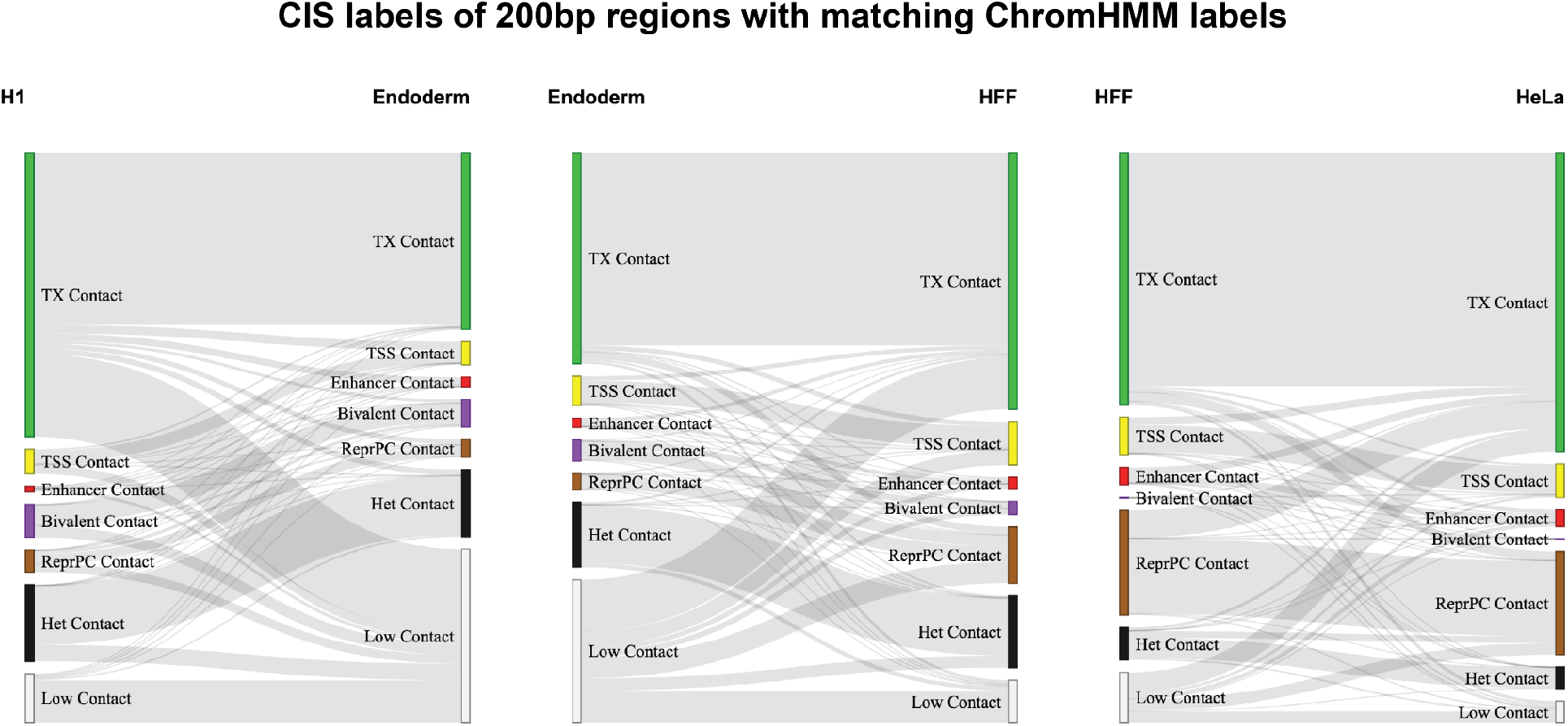
CIS change of regions with a constant ChromHMM state between cell types. For a window size of 200bp, CIS change is shown for regions with matching ChromHMM chromatin states between cell types.

**Figure S4.**
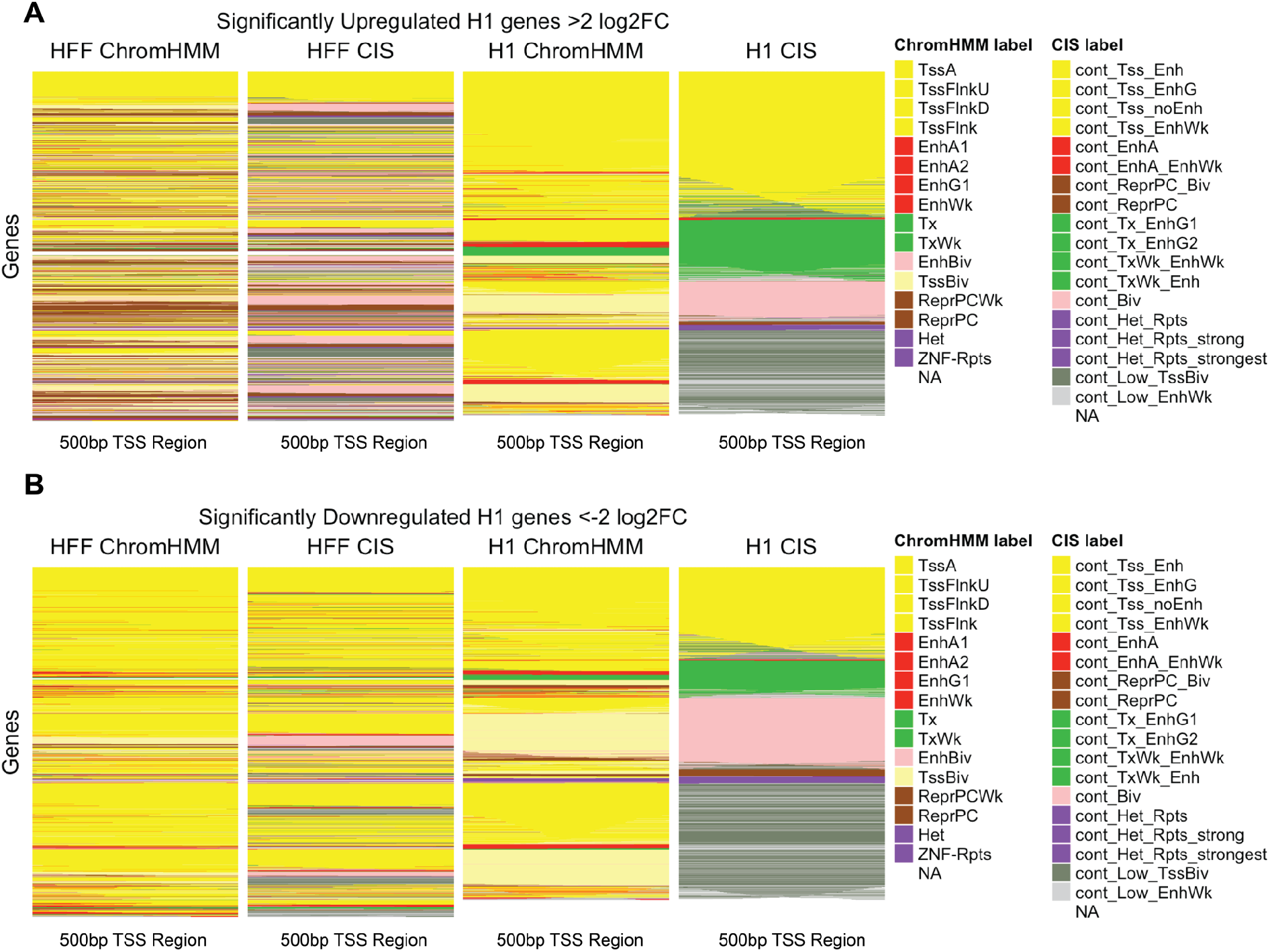
CIS and chromatin states for differentially expressed genes. Genes with adjusted p-value less than 1e-10 and with an H1 vs HFF log2 fold change >2 for upregulated genes (A) and <-2 for downregulated genes (B) were selected between H1 and HFF cell types. CIS and chromatin states are shown for each gene’s 500bp TSS region. The genes (heatmap rows) are ordered by proportion of clusters in order of H1 CIS, H1 chromatin state, HFF CIS, then HFF chromatin state.

